# The RgaS-RgaR two-component system promotes *Clostridioides difficile* sporulation through a small RNA and the Agr1 system

**DOI:** 10.1101/2023.06.26.546640

**Authors:** Adrianne N. Edwards, Shonna M. McBride

## Abstract

The ability to form a dormant spore is essential for the survival of the anaerobic, gastrointestinal pathogen *Clostridioides difficile* outside of the mammalian gastrointestinal tract. The initiation of sporulation is governed by the master regulator of sporulation, Spo0A, which is activated by phosphorylation. Multiple sporulation factors control Spo0A phosphorylation; however, this regulatory pathway is not well defined in *C. difficile*. We discovered that RgaS and RgaR, a conserved orphan histidine kinase and orphan response regulator, function together as a cognate two-component regulatory system to directly activate transcription of several genes. One of these targets, *agrB1D1*, encodes gene products that synthesize and export a small quorum-sensing peptide, AgrD1, which positively influences expression of early sporulation genes. Another target, a small regulatory RNA now known as SrsR, impacts later stages of sporulation through an unknown regulatory mechanism(s). Unlike Agr systems in many organisms, AgrD1 does not activate the RgaS-RgaR two-component system, and thus, is not responsible for autoregulating its own production. Altogether, we demonstrate that *C. difficile* utilizes a conserved two-component system that is uncoupled from quorum-sensing to promote sporulation through two distinct regulatory pathways.

**AUTHOR SUMMARY:** The formation of an inactive spore by the anaerobic gastrointestinal pathogen, *Clostridioides difficile*, is required for its survival outside of the mammalian host. The sporulation process is induced by the regulator, Spo0A; yet, how Spo0A is activated in *C. difficile* remains unknown. To address this question, we investigated potential activators of Spo0A. Here, we demonstrate that the sensor RgaS activates sporulation, but not by direct activation of Spo0A. Instead, RgaS activates the response regulator, RgaR, which in turn activates transcription of several genes. We found two direct RgaS-RgaR targets independently promote sporulation: *agrB1D1*, encoding a quorum-sensing peptide, AgrD1, and *srsR*, encoding a small regulatory RNA. Unlike most other characterized Agr systems, the AgrD1 peptide does not affect RgaS-RgaR activity, indicating that AgrD1 does not activate its own production through RgaS-RgaR. Altogether, the RgaS-RgaR regulon functions at multiple points within the sporulation pathway to tightly control *C. difficile* spore formation.

## INTRODUCTION

*Clostridioides difficile* is an anaerobic gastrointestinal pathogen that causes severe diarrheal disease (1). The vegetative cells of this strict anaerobe undergo a dramatic morphogenesis within the mammalian gastrointestinal tract to form dormant, highly resistant endospores. This infectious spore enables long-term survival outside of the host and facilitates transmission to new hosts (2). The initiation of spore formation requires the activation of the highly conserved transcriptional regulator, Spo0A, which is encoded in all endospore-forming bacteria (2–4). Spo0A DNA-binding activity is controlled by the phosphorylation of a conserved aspartate residue (5,6), allowing phosphorylated Spo0A (Spo0A∼P) to directly activate early sporulation-specific gene expression and triggering the entry into sporulation (5–8).

In other well-studied spore formers, most notably the soil dweller *Bacillus subtilis*, Spo0A phosphorylation is tightly controlled by an expansive regulatory network comprised of kinases, phosphatases, phosphotransferases, and additional regulators (9). *B. subtilis* Spo0A phosphorylation is achieved via a phosphorelay that begins with sensor histidine kinases that autophosphorylate and transmit a phosphoryl group through a response regulator (Spo0F) and a phosphotransferase (Spo0B) to Spo0A. Many of the key regulatory proteins that govern Spo0A activation in *B. subtilis* are not well conserved or are absent in *C. difficile* genome (10–13), suggesting that *C. difficile* utilizes unique factors and regulatory pathways to initiate spore formation. Three *C. difficile* phosphotransfer proteins (PtpA, PtpB, and PtpC), originally annotated as orphan histidine kinases, appear to inactivate Spo0A under the conditions tested (14,15). Thus, the primary driver of *C. difficile* Spo0A phosphorylation and activation is unknown.

Environmental cues and nutrient availability influence *C. difficile* sporulation initiation in the host gastrointestinal tract (16–20). Additionally, bacterial global signaling systems play a significant role in the decision to initiate sporulation. The intracellular nucleotide second messenger, c-di-GMP, is an inhibitor of early sporulation in *C. difficile* (21). In other spore formers, quorum-sensing systems are well established for regulating sporulation initiation (22,23). The RRNPP family of multifunctional proteins in *B. subtilis* and *Bacillus cereus* sp. directly bind small quorum-sensing peptides to regulate their activity, which impacts Spo0A phosphorylation (24,25). The *C. difficile* RRNPP ortholog, RstA, positively influences spore formation by directly binding an inhibitor of sporulation (26,27). However, RstA differs from most members of the RRNPP family in that its cognate quorum-sensing peptide is not adjacently encoded on the genome and has yet to be discovered (26), providing additional evidence that *C. difficile* employs distinct regulatory pathways to initiate sporulation.

Another conserved quorum sensing system implicated in spore formation is the accessory gene regulator (Agr) system, which utilizes an extracellular, cyclic autoinducing peptide (also generically known as the AIP) as the quorum-sensing signal. The prototypical Agr system encompasses the autoinducing peptide (AgrD), a transmembrane protease (AgrB) that processes and exports AgrD, a sensor histidine kinase (AgrC), which directly senses extracellular AgrD, and the cognate response regulator (AgrA) (28). Prior studies in *Clostridium botulinum*, *Clostridium sporogenes*, and *Clostridium perfringens* demonstrated that disruption or knockdown of the *agrBD* genes resulted in decreased spore formation (29,30). The majority of *C. difficile* genomes encode an *agrB1D1* operon with no apparent cognate *agrA1* or *agrC1* present (29,31–33). Recent work revealed that the *agrB1D1* locus positively impacts early sporulation gene expression and spore formation (34); however, the identity of the AgrD1 receptor remains elusive.

*C. difficile* encodes an orphan histidine kinase (CD0576; herein, RgaS) that is annotated as an ArgC-like sensor histidine kinase. Because the sporulation factor(s) that activate Spo0A are unclear, we asked if RgaS plays a role in *C. difficile* sporulation initiation. Our results reveal that RgaS signals through an orphan response regulator, RgaR, to promote *C. difficile* spore formation. We demonstrate that the conserved residues required for phosphoryl group transfer in both RgaS and RgaR are necessary for their function as a cognate two-component system. Further, RgaSR directly activate transcription of several loci, including *agrB1D1* and a small regulatory RNA, *n00620* (herein referred to as *srsR*, for <u>s</u>mall <u>R</u>NA <u>s</u>porulation <u>R</u>egulator). We establish that RgaSR influences spore formation through direct regulation of both *agrB1D1* and *srsR* transcription. Additionally, we show that the effects of AgrD1 and *srsR* are independent and impact different stages of sporulation. Unlike prototypical Agr systems, RgaS does not respond to the AgrD1 peptide, indicating that AgrB1D1 has adapted distinct functions that are independent of the RgaSR two-component system. Finally, we present evidence that RgaSR promotion of spore formation through AgrD1 and *srsR* is functionally conserved in diverse *C. difficile* strains.

## RESULTS

### The orphan histidine kinase RgaS and the RgaR response regulator promote *C. difficile* sporulation

The master response regulator of sporulation, Spo0A, is activated by phosphorylation. Multiple kinases activate sporulation in other spore-forming organisms; however, these factors are not conserved in *C. difficile*. Although three phosphotransfer proteins are proposed to act as phosphatases to inhibit Spo0A activity (15), a factor that directly phosphorylates Spo0A has not been discovered (13). We sought to determine if other encoded kinases could activate Spo0A to drive sporulation. We identified an additional, putative orphan histidine kinase within the *C. difficile* genome, *rgaS* (*CD0576*), that has limited similarity to the other sporulation-associated phosphotransfer proteins. To examine whether RgaS contributes to *C. difficile* sporulation, we first utilized CRISPR interference (CRISPRi) to directly repress *rgaS* gene expression. As the addition of xylose to sporulation medium reduces sporulation frequency (15), we modified the *C. difficile* CRISPRi plasmid (35) to drive expression of the nuclease-deactivated version of the *dCas9* gene with the nisin-inducible *cprA* promoter. A scrambled sgRNA (sgRNA-*neg*) was included as a control. We assessed the sporulation frequency of 630Δ*erm* expressing either sgRNA-*neg* or sgRNA-*rgaS* by determining the ratio of spores to vegetative cells after 24 h of growth on 70:30 sporulation agar, with and without nisin. Expressing the anti-*rgaS* target resulted in a ∼6- fold decrease in spore formation compared to the control strain (**Fig. 1A**). To ensure that *rgaS* was directly targeted, *rgaS* transcripts were measured by quantitative reverse- transcription PCR (qRT-PCR), and as expected, *rgaS* transcripts were decreased ∼25- fold in the presence of nisin (**Fig. S1A**). This data suggests that RgaS positively impacts

**Figure 1.**
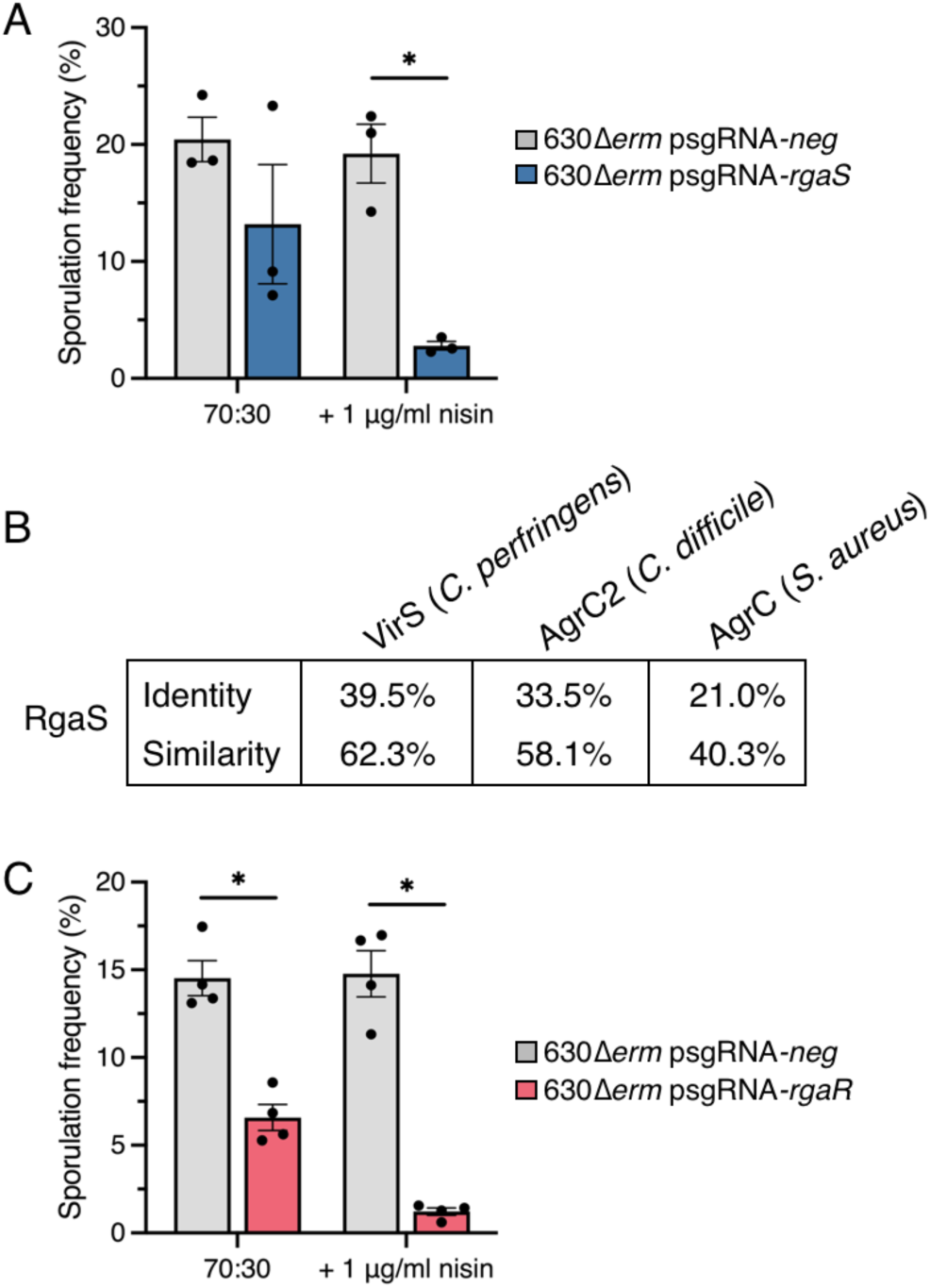
CRISPRi knockdown of *rgaS* (*CD0576*) and *rgaR* (*CD3255*) results in decreased sporulation frequency. (**A, C**) Ethanol-resistant spore formation at H_24_ in 630Δ*erm* psgRNA-*neg* (MC2065), 630Δ*erm* psgRNA-*rgaS* (MC2066), and 630Δ*erm* psgRNA-*rgaR* (MC2227) grown on 70:30 agar supplemented with 2 μg/ml thiamphenicol and 1 μg/m nisin, as indicated. The means and standard deviation of three independent biological replicates (A) or standard error of the means of four independent biological replicates (C) are shown. *, P < 0.01 by a Student’s *t-*test. (**B**) Percent identity and similarity of *Clostridium perfringens* VirS, *C. difficile* AgrC2 (CDR20291_3188), and *Staphylococcus aureus* AgrC to the histidine kinase RgaS in *C. difficile* 630.

RgaS possesses a histidine kinase-like ATPase domain that is similar to the AgrC histidine kinase of *Staphylococcus aureus* (**Fig. 1B**). Additional AgrC-like kinases have been identified in *Clostridia*, including the *Clostridium perfringens* VirS (36) and the *C. difficile* AgrC2, which is encoded only in the epidemic ribotype 027 strains (37). Recent work established that *C. perfringens* VirS, which directly phosphorylates its cognate response regulator, VirR (38), responds to the Agr quorum sensing peptide produced by *C. perfringens* to activate VirR-dependent gene expression (39). RgaS shares high similarity and identity with *C. perfringens* VirS and *C. difficile* R20291 AgrC2 (**Fig. 1B**), suggesting that RgaS functions as a histidine kinase that directly senses an extracellular quorum sensing peptide.

We hypothesized that if RgaS is a VirS-like histidine kinase, then its cognate response regulator would be similar to *C. perfringens* VirR. We scanned the *C. difficile* 630 genome for orphan response regulators and located two VirR orthologs: CD1089 (RgbR) and CD3255 (RgaR). A prior investigation showed that overexpression of *rgaR* in a *C. perfringens virR* mutant partially complemented toxin production, suggesting that RgaR functions as a VirR ortholog (40). Further, they discovered that RgaR directly activated transcription of three *C. difficile* operons, including the partial *agrB1D1* locus (40). To determine whether RgaR affects *C. difficile* sporulation, we knocked down *rgaR* gene expression by CRISPRi. *C. difficile* sporulation was significantly decreased (∼12- fold) in the 630Δ*erm* strain expressing sgRNA-*rgaR* compared to the parental control, indicating that RgaR positively influences spore formation (**Fig. 1C**). *rgaR* transcript levels were decreased ∼4-fold (**Fig. S1B)**, confirming that the *rgaR* locus was directly targeted by CRISPRi.

To further explore the impact of RgaS and RgaR on *C. difficile* sporulation, we generated *rgaS* and *rgaR* deletion mutants using allele-coupled exchange (41) (see Methods; **Fig. S2**). Sporulation phenotypes of the mutants were assessed after 24 h growth on sporulation agar. The *rgaS* and *rgaR* single mutants sporulated at lower frequencies (∼7%) than the 630Δ*erm* parent (∼21%; **Fig. 2A**), corroborating the CRISPRi knockdown results. These sporulation phenotypes were also confirmed by phase contrast microscopy (**Fig. 2B**), as fewer phase bright spores are visible in the *rgaS* and *rgaR* mutants. We also created a double *rgaS rgaR* mutant and found that the double mutant exhibited similar sporulation frequencies as the single *rgaS* and *rgaR* mutants (**Fig. 2C**), suggesting that RgaS and RgaR function in the same regulatory pathway to affect sporulation.

**Figure 2.**
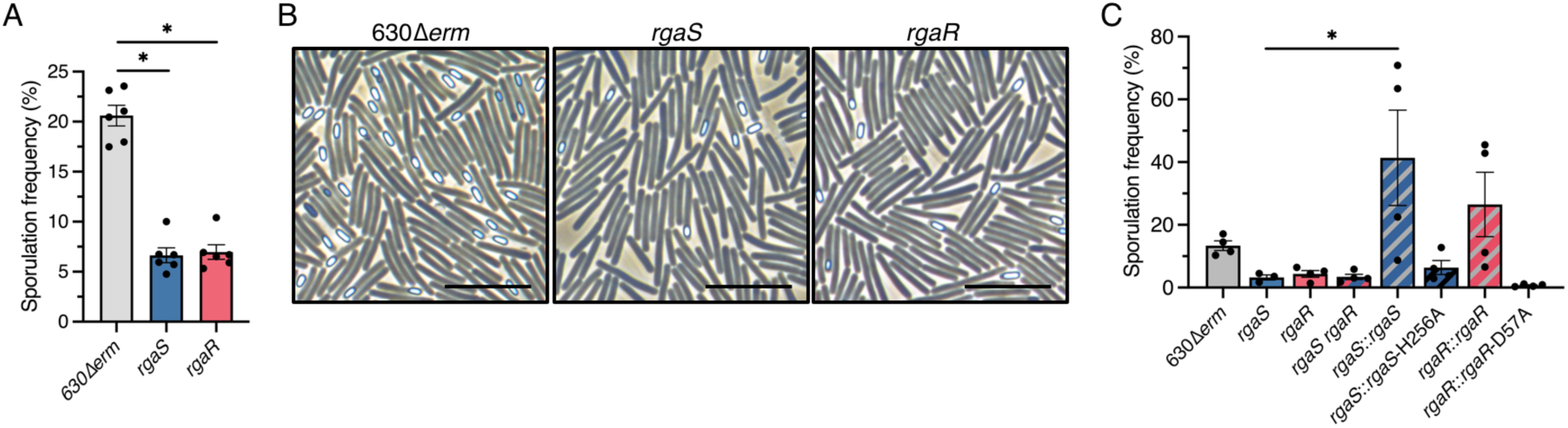
*C. difficile rgaS* and *rgaR* mutants exhibit low sporulation frequencies and require their conserved histidine or aspartate residue for function. (**A**) Ethanol-resistant spore formation and (**B**) representative phase-contrast micrographs at H_24_ in 630Δ*erm*, 630Δ*erm* Δ*rgaS* (MC2228), and 630Δ*erm* Δ*rgaR* (MC2229) grown on 70:30 agar. Scale bar denotes 10 μm. (C) Ethanol-resistant spore formation at H_24_ of 630Δ*erm*, 630Δ*erm* Δ*rgaS* (MC2228), 630Δ*erm* Δ*rgaR* (MC2229), 630Δ*erm* Δ*rgaS* Δ*rgaR* (MC2236), 630Δ*erm* Δ*rgaS* Tn*916*::*rgaS* (MC2278), 630Δ*erm* Δ*rgaS* Tn*916*::*rgaS-*H256A (MC2318), 630Δ*erm* Δ*rgaR* Tn*916*::*rgaR* (MC2319), and 630Δ*erm* Δ*rgaR* Tn*916*::*rgaR-*D57A (MC2320) grown on 70:30 agar. The means and standard error of the means of at least four independent biological replicates are shown. *, P < 0.01 by a one-way ANOVA followed by Dunnett’s multiple comparisons test (A) or Tukey’s multiple comparisons test (B).

To ensure that the decreased sporulation frequencies of the *rgaS* and *rgaR* mutants are solely the result of the deleted genes, we complemented these mutants by expressing the wild-type *rgaS* and *rgaR* alleles from their native promoters on the chromosome via the Tn*916* conjugative transposon. Expression of the *rgaS* and *rgaR* alleles in their respective mutants restored sporulation frequencies to greater than wild-type levels (**Fig. 2C**).

Our data suggest that RgaS and RgaR function in the same regulatory pathway as a two-component regulatory system (TCS), where the RgaS histidine kinase undergoes autophosphorylation at the conserved histidine residue and transfers the phosphoryl group to the conserved aspartate residue of the RgaR response regulator (42). To determine whether the predicted functional residues of RgaS and RgaR are essential for their activity, we replaced the conserved histidine residue of RgaS and the conserved aspartate residue of RgaR with alanine, using site-directed mutagenesis, and expressed these alleles in their respective mutants via the Tn*916* conjugative transposon. Neither the *rgaS*-H256A nor the *rgaR*-D57A allele complemented the low sporulation phenotypes of the respective mutants (**Fig. 2C**), demonstrating that these residues are required for RgaS and RgaR to promote sporulation.

### Late-stage sporulation is downregulated in the *rgaR* mutant

To better understand how the RgaS-RgaR TCS promote sporulation, we performed RNA-seq on the *rgaR* mutant during growth on sporulation agar (H_12_) to identify differentially expressed genes during sporulation. The development of spores requires the activation of Spo0A followed by the sequential activation of the sporulation sigma factors, SigF, SigE, SigG, and SigK (10,43). RNA-seq analysis revealed that only a few Spo0A- or SigF-dependent transcripts were significantly decreased in the *rgaR* mutant (∼22% of the Spo0A regulon and ∼33% of the SigF regulon; **Table 1**), suggesting that RgaR does not strongly influence the initiation of sporulation. However, ∼84% of the SigE regulon, 94% of the SigG, and 97% of the SigK regulons were significantly downregulated in the *rgaR* mutant (**Table 1**), revealing that RgaR promotes the progression to later stages of spore formation.

**Table 1.** Late-stage sporulation genes are significantly downregulated in the *rgaR* mutant.

| Regulon<sup>a</sup> | Upregulated genes | Unaffected genes | Downregulated genes | Total genes in regulon<sup>b</sup> | Percent of genes downregulated<sup>c</sup> |
| --- | --- | --- | --- | --- | --- |
| Spo0A-dependent | 12 | 56 | 19 | 87 | 22% |
| SigF-dependent | 0 | 14 | 7 | 21 | 33% |
| SigE-dependent | 0 | 17 | 89 | 106 | 84% |
| SigG-dependent | 0 | 2 | 31 | 33 | 94% |
| SigK-dependent | 0 | 1 | 29 | 30 | 97% |

<sup>a</sup> The sporulation-associated sigma factor regulons are based on those reported in (7).
<sup>b</sup> Genes whose expression were dependent on multiple sporulation-specific sigma factors were excluded from this analysis.
<sup>c</sup> These values are based on fold changes from the RNA-seq analyses of 630Δ*erm* Δ*rgaR* (MC2229) compared to the 630Δ*erm* parent strain grown on 70:30 agar at H<sub>12</sub>.

The RNA-seq results also revealed that expression of the toxin genes *tcdA* and *tcdB* were largely unaffected in the *rgaR* mutant, with a modest ∼2-fold decrease in *tcdB* transcripts (**Table S1**). We further assessed toxin production by measuring the total TcdA and TcdB toxin present in the supernatants of the *rgaS* and *rgaR* mutants. Toxin levels were unchanged between the *rgaS* mutant, the *rgaR* mutant, and the parent strain (**Fig. S3**). Altogether, these data confirm that the RgaSR TCS does not substantially impact toxin production in the conditions tested and suggests that the effects of RgaSR on sporulation regulation are independent of toxin regulation.

### RgaS and RgaR have the same effects on gene expression

Prior work revealed that RgaR directly binds and activates transcription from three promoters that contain highly conserved VirR-like consensus binding sites (40). These direct RgaR targets include the promoters for *agrB1D1*, the hypothetical factor *CD2098*, and *CD0587-CD0588*, a two-gene operon encoding two hypothetical proteins. Bioinformatical mining uncovered two additional VirR-like binding sites located in intergenic regions within the *C. difficile* genome (40), which have since been annotated. One VirR-like binding site is located upstream of a small ORF encoding a hypothetical protein, CD15111, and the second site lies immediately upstream of a predicted small regulatory RNA we have dubbed SrsR (**Fig. 3A**). We next asked if these direct RgaR targets are similarly regulated in the *rgaS* mutant. Using qRT-PCR, we assessed transcript levels of RgaR-regulated genes in sporulating cells and observed the same decrease in transcript levels for the *rgaS* and *rgaR* mutants (**Fig. 3B**). Altogether, these data strongly support that RgaS is the cognate histidine kinase of the RgaR response regulator and that these proteins comprise a functional two-component regulatory system.

**Figure 3.**
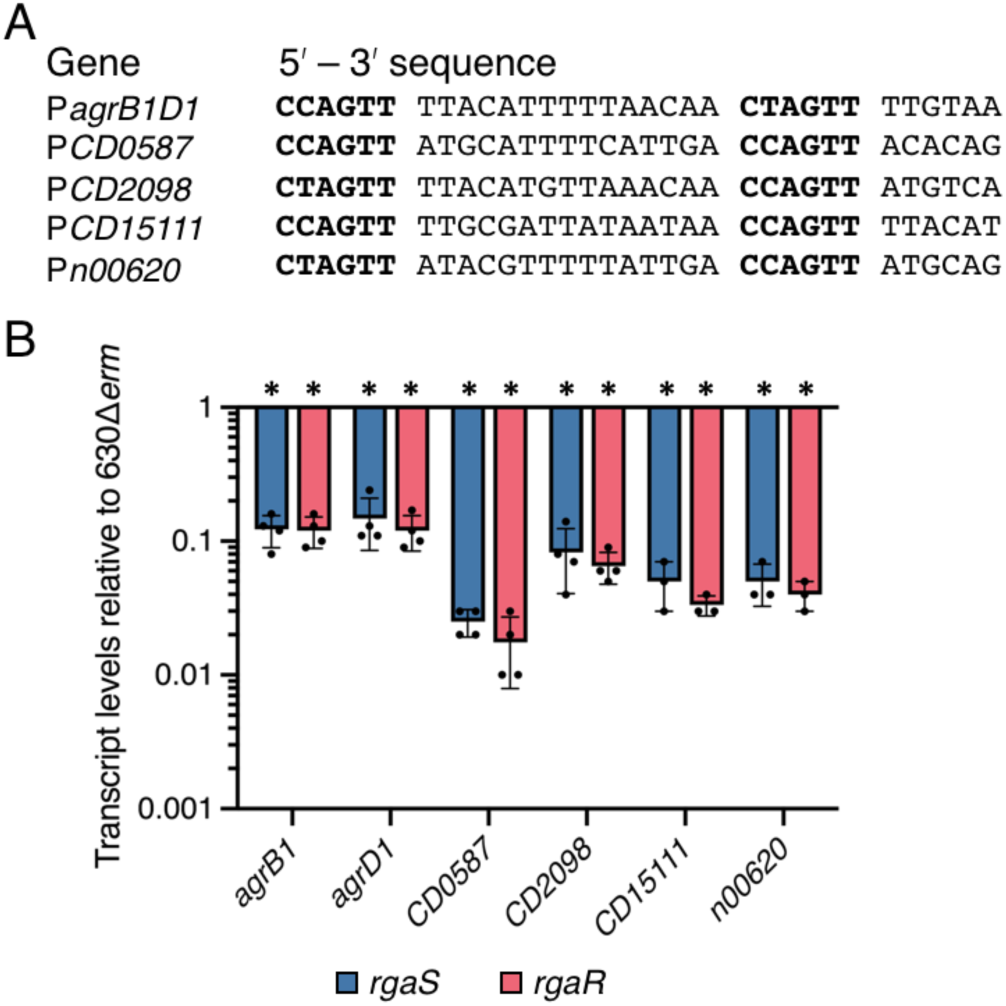
RgaR-dependent gene expression is similarly decreased in both the *rgaS* and *rgaR* mutants. (**A**) Alignment of the putative RgaR binding sites with the promoters of confirmed and predicted direct RgaR targets. The direct repeats are in bold. (**B**) qRT-PCR analysis of RgaR-dependent transcripts at H_12_ in 630Δ*erm*, 630Δ*erm* Δ*rgaS* (MC2228), and 630Δ*erm* Δ*rgaR* (MC2229) grown on 70:30 agar. The means and standard error of the means of at least three independent biological replicates are shown. *, P < 0.0001 by a one-way ANOVA followed by Dunnett’s multiple comparisons test.

### RgaR regulates sporulation through the small regulatory RNA, SrsR

To determine which direct RgaR target is responsible for the low sporulation phenotype in the *rgaS* and *rgaR* mutants, we used CRISPRi to individually knockdown transcription of *CD0587*, *CD2098*, *CD15111*, and *srsR,* and assessed the impacts on sporulation. Knockdown of *CD0587*, *CD2098*, or *CD15111* did not alter sporulation frequency compared to the control (**Fig. 4A**). However, the knockdown of *srsR* significantly increased spore formation (**Fig. 4A**), suggesting that the RgaS and RgaR influence sporulation frequency through the SrsR small regulatory RNA. As before, we confirmed direct targeting of transcription by CRISPRi using qRT-PCR (**Fig. S4**).

**Figure 4.**
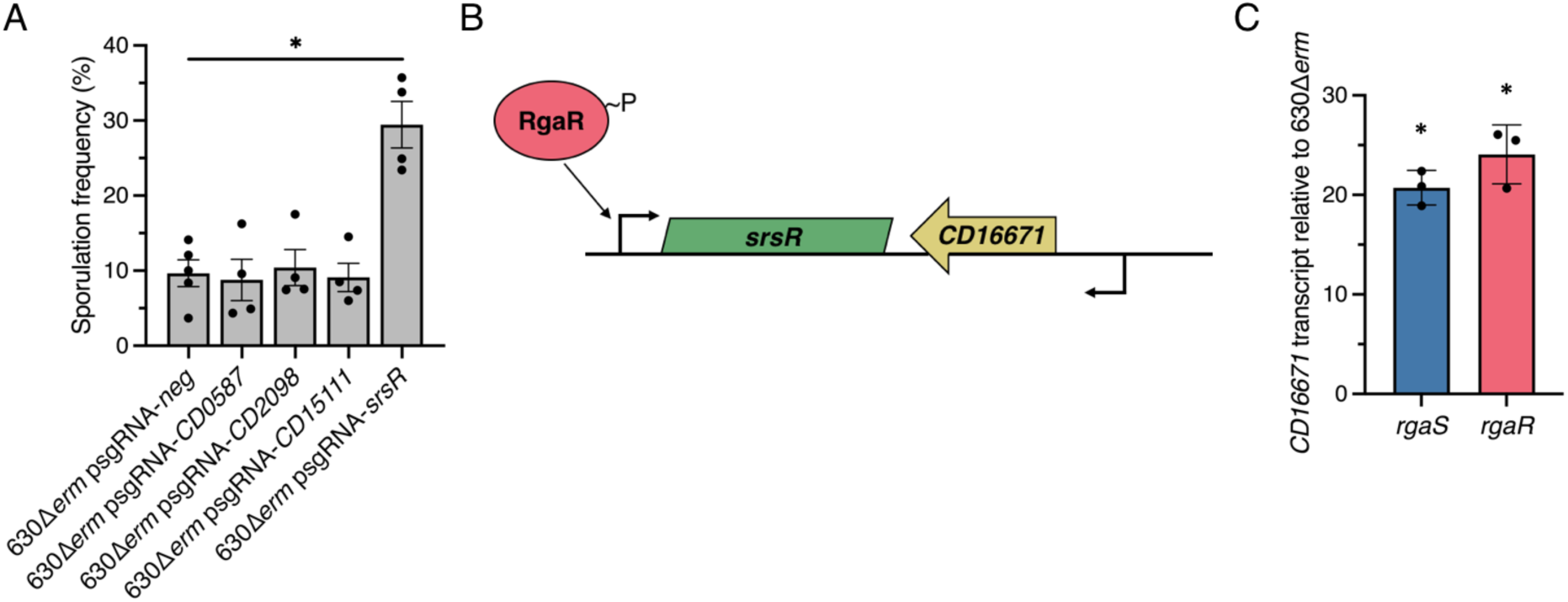
CRISPRi knockdown of RgaR-dependent genes reveal that RgaR impacts sporulation through the regulatory sRNA, SrsR (encoded by *CD630_n00620*). (**A**) Ethanol-resistant spore formation of 630Δ*erm* expressing p*sgRNA-neg* (MC2065) or CRISPRi targets for *CD0587* (MC2269), *CD2098* (MC2270), *CD15111* (MC2271), or *srsR* (MC2272) grown on 70:30 agar supplemented with 2 μg/ml thiamphenicol and 1 μg/m nisin at H_24_. The means and standard errors of the means for four biological replicates are shown. (**B**) Genomic context of the sRNA, *srsR*, in relation to the small ORF, *CD16671*. (**C**) qRT-PCR analysis of *CD16671* in 630Δ*erm*, 630Δ*erm* Δ*rgaS* (MC2228), and 630Δ*erm* Δ*rgaR* (MC2229) grown on 70:30 agar at H_12_. The means and standard deviations of three independent biological replicates are shown. *, *P* < 0.01 by one-way ANOVA followed by Dunnett’s multiple comparisons test.

SrsR was originally identified as a non-coding intergenic RNA that was predicted to act as an antisense target for the neighboring ORF, CD16671 (44–46). Subsequent studies discovered that SrsR copurifies with Hfq, an RNA-binding protein that mediates post-transcriptional RNA interactions. We hypothesized that SrsR functions as an antisense RNA that targets the *CD16671* transcript for Hfq-dependent turnover (**Fig. 4B**). We assessed *CD16671* transcript levels in the *rgaS* and *rgaR* mutants and found that the *CD16671* transcript is significantly increased in these mutants (**Fig. 4C**). These results suggest that SrsR may promote sporulation by inhibiting accumulation of the *CD16671* transcript and subsequent protein product.

To test this hypothesis, we generated deletion mutants of either the entire *srsR- CD16671* locus or only *CD16671* (see Methods; **Fig. S5A and S5B**) and assessed their sporulation phenotypes. Like the *srsR* knockdown strain, deletion of *srsR* resulted in increased sporulation frequency (∼2.7-fold) compared to the parent strain (**Fig. 5A**). However, deletion of *CD16671* did not impact sporulation frequency, implying that *CD16671* is not involved in sporulation regulation. Thus, SrsR must have an unidentified target(s) that mediates its effects on sporulation.

**Figure 5.**
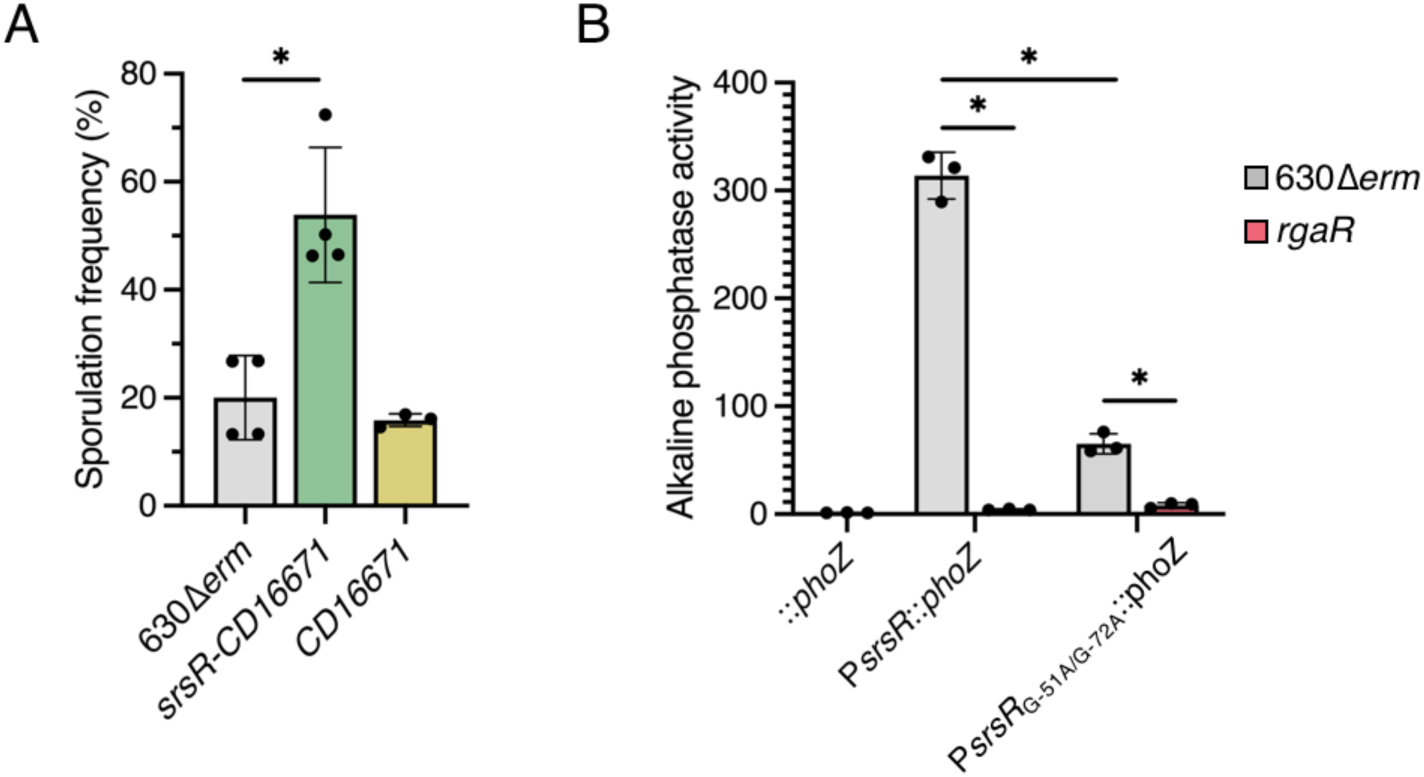
The RgaR-regulated sRNA, SrsR, negatively affects *C. difficile* spore formation. (**A**) Ethanol-resistant spore formation of 630Δ*erm,* 630Δ*erm* Δ*srsR-CD16671* (MC2351), and 630Δ*erm* Δ*CD16671* (MC2363) grown on 70:30 agar at H_24_. (**B**) Alkaline phosphatase activity of either the wildtype P*srsR::phoZ* or a site-directed mutagenized P*srsR*_G-51A/G-72A_*::phoZ* reporter fusion expressed from a plasmid in 630Δ*erm* (MC2329 and MC2357, respectively) or 630Δ*erm* Δ*rgaR* (MC2330 and MC2358, respectively). The promoterless *phoZ* reporter carried by 630Δ*erm* (MC448) was included as a negative control. The means and standard deviations for at least three biological replicates are shown. *, *P* < 0.01 by one-way ANOVA followed by Dunnett’s multiple comparisons test (A) or Tukey’s multiple comparisons test (B).

The *srsR-CD16671* mutant was confirmed via whole genome sequencing; However, despite numerous attempts with various constructs and approaches, we were unable to complement the *srsR-CD16671* mutant (also see Discussion). To ensure that the transcript(s) generated from the *srsR-CD16671* locus were included in our complementation constructs, we analyzed the cDNA synthesized from this region in the parent strain (**Fig. S5C, D**). We were able to amplify a small product corresponding to the predicted 278 nt sRNA, as well as a larger product extending through the entire coding region of *CD16671* (**Fig. S5C**). But, we could not amplify a PCR product extending beyond the predicted *CD16671* promoter, indicating that the *srsR* transcript terminates within the *CD16671* promoter region (**Fig. S5D**). These data corroborate the previously mapped transcription termination site for *srsR*, slightly beyond the *CD16671* ORF (528 nt) (45). In all, the regulatory portion of SrsR is not well defined and additional studies are needed to delineate the molecular role SrsR plays in *C. difficile* spore formation.

### AgrD1 does not signal through the RgaS-RgaR two component system

The prototypical Agr signaling systems form a regulatory feedback loop, in which the quorum peptide signal (AgrD) activates the sensor kinase to initiate Agr-dependent regulation. As RgaS-RgaR directly activates *agrB1D1* transcription and a previously published *agrB1D1* mutant exhibits lower sporulation frequency (34), we hypothesized that the AgrD1 quorum-sensing peptide is recognized by RgaS to activate RgaR-dependent gene expression. To test this, we created an *agrB1D1* deletion mutant by allelic exchange (see Methods; **Fig. S6A**). The *agrB1D1* mutant made fewer spores, similar to the *rgaS* and *rgaR* mutants (**Fig. 6A**), and expressing *agrB1D1* from an inducible promoter restored the sporulation frequency to greater than wild-type levels.

**Figure 6.**
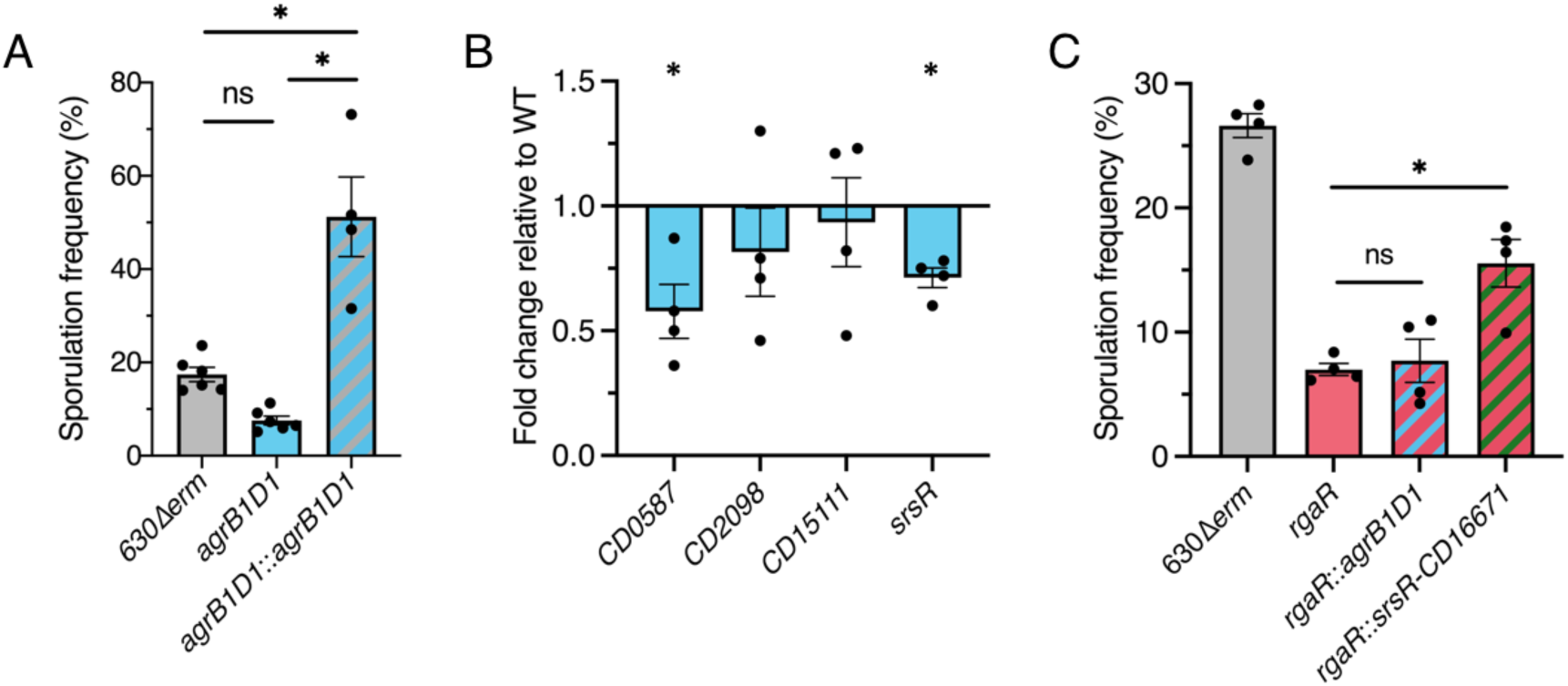
AgrB1D1 does not activate RgaR-dependent gene expression and promotes sporulation upstream of SrsR. (**A**) Ethanol-resistant spore formation of 630Δ*erm*, 630Δ*erm* Δ*agrB1D1* (MC2433), and 630Δ*erm* Δ*agrB1D1* Tn*916*::P*cprA- agrB1D1* (MC2524) grown on 70:30 agar at H_24_. (**B**) Transcript levels of RgaR- dependent genes, *CD0587*, *CD2098*, *CD15111*, and *srsR in* 630Δ*erm*, 630Δ*erm* Δ*agrB1D1* (MC2433), and 630Δ*erm* Δ*agrB1D1* Tn*916*::P*cprA-agrB1D1* (MC2524) grown on 70:30 agar at H_12_. (**C**) Ethanol-resistant spore formation of 630Δ*erm*, 630Δ*erm* Δ*rgaR* (MC2229), 630Δ*erm* Δ*rgaR* Tn*916*::P*cprA-agrB1D1* (MC2424), and 630Δ*erm* Δ*rgaR* Tn*916*::P*cprA-srsR-CD16671* (MC2425) grown on 70:30 agar at H_24_. The means and standard errors of the means for at least four biological replicates are shown. *, *P* < 0.01 by Student’s *t-*test (B) or an one-way ANOVA followed by Tukey’s multiple comparisons test (A, C).

We next asked whether the RgaS-RgaR-dependent gene expression was dependent on AgrD1. To answer this question, we assessed RgaR-dependent gene expression in the *agrB1D1* mutant. Surprisingly, qRT-PCR analysis revealed that transcript levels of *CD0587*, *CD2098*, *CD15111*, and *srsR* were not dramatically reduced in the *agrB1D1* mutant (**Fig. 6B**). Thus, RgaS is not the sensor for the AgrD1 peptide, as expression of RgaR-dependent genes is not affected by AgrD1. This unconnected decrease in sporulation for the *agrB1D1* mutant indicates that AgrB1D1 promotes sporulation through a mechanism that is independent of RgaSR.

Prior analyses of transcription in *agrB1*, *agrD1*, and *agrB1D1* mutants revealed that AgrD1 promotes transcription of early sporulation genes (34). In contrast, disruption of *rgaR* most significantly impacted expression of late stage sporulation genes (**Table 1**), implying that SrsR affects later stages of sporulation. As the *rgaR* mutant has significantly decreased transcript levels of both *agrB1D1* and *srsR* (**Fig. 3B**), we asked whether AgrD1 or SrsR promote different stages of sporulation by performing an epistasis study in the *rgaR* mutant. We overexpressed either the *agrB1D1* or the *srsR- CD16671* locus in the *rgaR* mutant and assessed sporulation frequency. We anticipated that if SrsR is epistatic to AgrD1 (that is, if SrsR functions downstream from AgrD1 in the sporulation regulatory pathway) that only *srsR* overexpression will restore sporulation in the *rgaR* mutant. As expected, overexpression of *agrB1D1* did not increase sporulation of the *rgaR* mutant; however, overexpression of *srsR-CD16671* in the *rgaR* mutant doubled spore formation (**Fig. 6C**), demonstrating that *srsR* is able to partially complement the low sporulation phenotype of the *rgaR* mutant. Since AgrD1 supports early sporulation, overexpression of *agrB1D1* cannot compensate for an *rgaR* mutant, as SrsR must be required for later sporulation.

### The RgaS-RgaR regulatory network is conserved in the epidemic R20291 background

As the *agrB1D1* locus is highly conserved in almost all sequenced *C. difficile* genomes, we asked whether the RgaS-RgaR system is present and functions similarly in the epidemic ribotype 027 strain, R20291. The R20291 genome encodes highly conserved RgaS and RgaR orthologs, *CDR20291_0503* and *CDR20291_3113*. Null *rgaS* and *rgaR* mutants were created in the R20291 background (see Methods; **Fig. S6B**). We assessed sporulation frequency and observed that the *rgaS* and *rgaR* mutants produced fewer spores (∼1.6-fold and ∼2.1-fold, respectively) than the R20291 parent strain (**Fig. 7A**). Transcriptional analysis of the direct RgaR targets, *agrB1D1*, the *CD2098* ortholog (*CDR20291_2005*), the *CD15111* ortholog (unannotated in R20291; herein referred to as *CD15111^*), and *srsR*, revealed significantly decreased transcript levels in the *rgaS* and *rgaR* mutants, indicating that the RgaS-RgaR orthologs in R20291 function similarly as in 630Δ*erm*.

**Figure 7.**
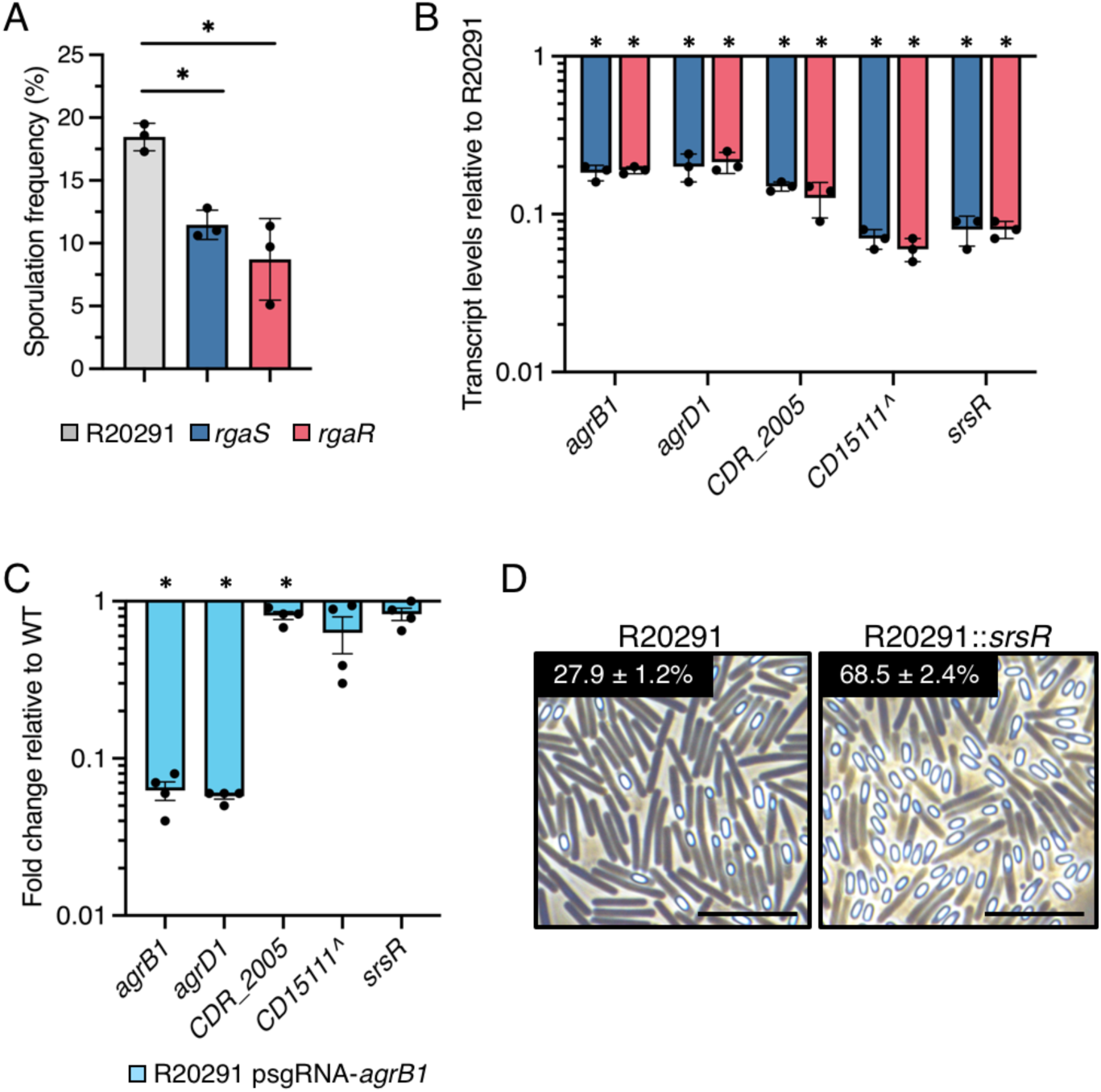
The RgaS-RgaR-AgrB1D1-SrsR regulatory network is conserved in R20291. (**A**) Ethanol-resistant spore formation at H_24_ and (**B**) qRT-PCR analyses of orthologous RgaR-dependent transcripts at H_12_ in R20291, R20291 Δ*rgaS* (MC3278), and R20291 Δ*rgaR* (MC3279) grown on 70:30 agar. The means and standard deviations of three independent biological replicates are shown. (**C**) qRT-PCR analyses of orthologous RgaR-dependent transcripts in R20291 psgRNA-*neg* (MC2237) and R20291 psgRNA-*agrB1* (MC2534) grown on 70:30 agar at H_12_. (**D**) Representative phase-contrast micrographs of R20291 and R20291 Tn*916*::P*cprA-srsR-CD16671* (MC2533) grown on 70:30 agar at H_24_. Numbers below represent the means and standard error of the means of four independent biological replicates from ethanol-resistant sporulation assays at H_24_. Scale bar denotes 10 μm. ^, The *CD15111* locus in 630Δ*erm* is not annotated in the R20291 genome used for this study, so the 630Δ*erm* locus tag is used as a placeholder. *, *P* < 0.01 by one-way ANOVA followed by Dunnett’s multiple comparisons test (A) or a student’s *t-*test (B-D).

To further evaluate the RgaS-RgaR and AgrD1 regulatory circuitry in the R20291 background, we next asked whether the R20291 RgaS-RgaR system was responsive to the AgrD1 quorum-sensing peptide. To do this, we utilized CRISPRi to knockdown *agrB1* transcription, which significantly decreased transcript levels of both *agrB1* and *agrD1* (**Fig. 7C**). We then assessed transcript levels of the direct RgaR targets, *CDR20291_2005*, *CD15111^*, and *srsR*, which remained unchanged in the *agrB1D1* knockdown (**Fig. 7C**). These data show that the RgaS-RgaR system also does not respond to AgrD1 in the R20291 background, indicating that this uncoupled signaling circuitry is conserved in diverse *C. difficile* strains. Finally, to determine whether the regulatory sRNA SrsR also impacts sporulation in R20291, we overexpressed the *srsR- CD16671* locus. Sporulation frequency in the *srsR-CD16671* overexpression strain was increased 2.5-fold (**Fig. 7D**), demonstrating that SrsR significantly impacts spore formation in R20291 as well. Altogether, these data suggest that the unusual RgaS-RgaR, AgrD1, and SrsR regulatory circuitry uncovered in 630Δ*erm* are conserved and functional in the R20291 background.

## DISCUSSION

Although the morphological changes throughout spore formation are conserved between clostridia and the well-studied bacilli, the regulatory pathways and molecular mechanisms that govern sporulation have significantly diverged. Further, the mechanism of Spo0A phosphorylation, which activates sporulation, is unclear in *C. difficile*. While searching for sporulation factors that facilitate *C. difficile* sporulation initiation, we uncovered a complex regulatory circuit that promotes sporulation through an orthologous accessory gene regulator (Agr) system. We discovered a non-contiguous, cognate two-component system, comprised of the transmembrane histidine kinase RgaS and the response regulator RgaR. RgaS-RgaR directly activate transcription of several loci, including the *agrB1D1* operon and the regulatory small RNA, *srsR*. We further demonstrated that the gene products of these loci, the quorum-sensing peptide, AgrD1, and the regulatory sRNA, SrsR, impact *C. difficile* spore formation at different stages of sporulation. Finally, our investigation revealed that AgrD1 does not activate the RgaS-RgaR two-component system, breaking the conventional positive feedback featured in characterized Agr systems.

To our knowledge, this is the first identification of an Agr-like two-component system that activates the transcription of the gene encoding the AIP precursor but does not serve as the receptor for the AIP. Classical Agr systems are activated through a positive feedback loop in which the histidine kinase (AgrC) autophosphorylates in response to the accumulation of AIP during high cell density conditions and subsequently transfers the phosphoryl group to the response regulator (AgrA), which in turn, activates transcription of target genes, including the genes that encode the Agr system components (47–50). Our findings reveal an unusual Agr system in *C. difficile* that is not autoregulatory. Previous work demonstrated that an *agrD1* mutant does not impact *agrB1* transcript levels (34), corroborating that the AgrD1 peptide does not influence its own transcription. *C. perfringens* encodes a single *agrB1D1* locus, and the VirS-VirR two component system is the AIP receptor (51,39); however, VirR does not activate *agrB1D1* transcription (52), also breaking the Agr autoregulatory loop. These examples suggest that clostridial Agr systems are not autoregulatory in general, although different mechanisms have evolved to unlink AIP accumulation to its synthesis. Considering that the *C. difficile* AgrD1 AIP is important for promoting spore formation independently of RgaS-RgaR, there must be another unidentified transmembrane sensor that serves as the AgrD1 receptor. Discovery of the *C. difficile* AgrD1 receptor will be important in elucidating the regulatory pathway through which AgrD1 promotes sporulation.

A striking feature of the RgaS-RgaR regulon is the direct transcriptional activation of a regulatory RNA. The regulon of Agr systems generally includes a gene encoding a small regulatory RNA. The most notable examples include the RNAIII regulatory RNA of *Staphylococcus aureus*, which is activated by AgrC-AgrA, and the VR-RNA, VirT, and VirU small regulatory RNAs in *C. perfringens*, which are activated by VirS-VirR (53–55). These regulatory RNAs function as global regulators in both *S. aureus* and *C. perfringens* by influencing transcript stability and translational efficiency of target mRNAs (56–60). Our discovery that RgaS-RgaR directly activates *srsR* transcription suggests that *C. difficile* also coordinately regulates multiple targets through a single regulatory sRNA.

The function of SrsR during *C. difficile* sporulation is difficult to discern because the sporulation phenotypes associated with *srsR* abundance appear internally inconsistent. Low levels of *srsR* transcript levels in the *rgaR* mutant (∼5% compared to the parent strain) led to a low sporulation frequency, while deletion of *srsR* resulted in hypersporulation. These conflicting sporulation phenotypes suggest that SrsR function varies with its abundance. The inability to complement the *srsR-CD16671* mutant may also be due to differences in the intracellular concentration and accumulation of SrsR. Several of the complementation constructs we tested contained the entire *srsR* transcript but did not restore *srsR* expression to wild-type levels (data not shown), further corroborating that the abundance of SrsR is important for its function. Additionally, a recent study employing Hfq RIL-seq identified a few direct sRNA/mRNA interactions with SrsR (61), indicating that SrsR has multiple direct targets. It is possible that SrsR exhibits higher affinity to a subset of targets and that more than one of these identified interactions is responsible for the different regulatory effects of SrsR on sporulation. These are not mutually exclusive hypotheses; future directions will focus on the molecular mechanisms utilized by SrsR to influence *C. difficile* sporulation.

Surprisingly, we found that our 630Δ*erm agrB1D1* mutants exhibited a variable sporulation phenotype that was selectable on plates. We observed the same phenomenon when *agrB1* was knocked down in the R20291 background (data not shown); however, this variation was not noted with the previously published *agrB1D1* mutant, which was also constructed in the 630Δ*erm* background. Since both low and high sporulation phenotypes can be differentiated on plates after passaging a colony exhibiting a single colony morphology, we propose that there is a phase variation event occurring with higher frequency in the *agrB1D1* mutants that influences sporulation. Phase variation is a common strategy used by *C. difficile* to exhibit significant phenotypic heterogeneity (62–68), and there is a possibility that the Agr1 system may regulate a recombinase or other regulatory factor that influences the phase variation of sporulation.

The components of the RgaS-RgaR-SrsR-AgrB1D1 regulatory circuitry are highly conserved in *C. difficile* genomes, suggesting these regulatory pathways are important in controlling earlier and later sporulation events across the species. The universal function of this system in *C. difficile* is further supported by data from the R20291 strain, which demonstrated conservation of the RgaS-RgaR regulon and sporulation phenotypes. In addition, bioinformatic analysis of the R20291 genome revealed the same RgaR targets, with no additional RgaR binding sites compared to the 630Δ*erm* strain.

Our working model of the Rga regulatory system (**Fig. 8**) begins with RgaS autophosphorylation and phosphoryl group transfer to RgaR, which then directly activates both *agrB1D1* and *srsR* transcription. The resulting pro-AgrD1 is processed and exported by AgrB1, accumulates extracellularly in high cell density conditions and promotes the initiation of sporulation through an unknown factor. The small regulatory RNA, SrsR, targets one or more sRNA/mRNA(s) to influence late stage sporulation. In all, the RgaS-RgaR regulon integrates at multiple points within the sporulation pathway to control *C. difficile* spore formation. Understanding when RgaSR is most active and which signals activate (or inhibit) this system will aid in understanding the environmental cues that promote *C. difficile* sporulation, permitting its survival in the environment and efficient transmission to new hosts.

**Figure 8.**
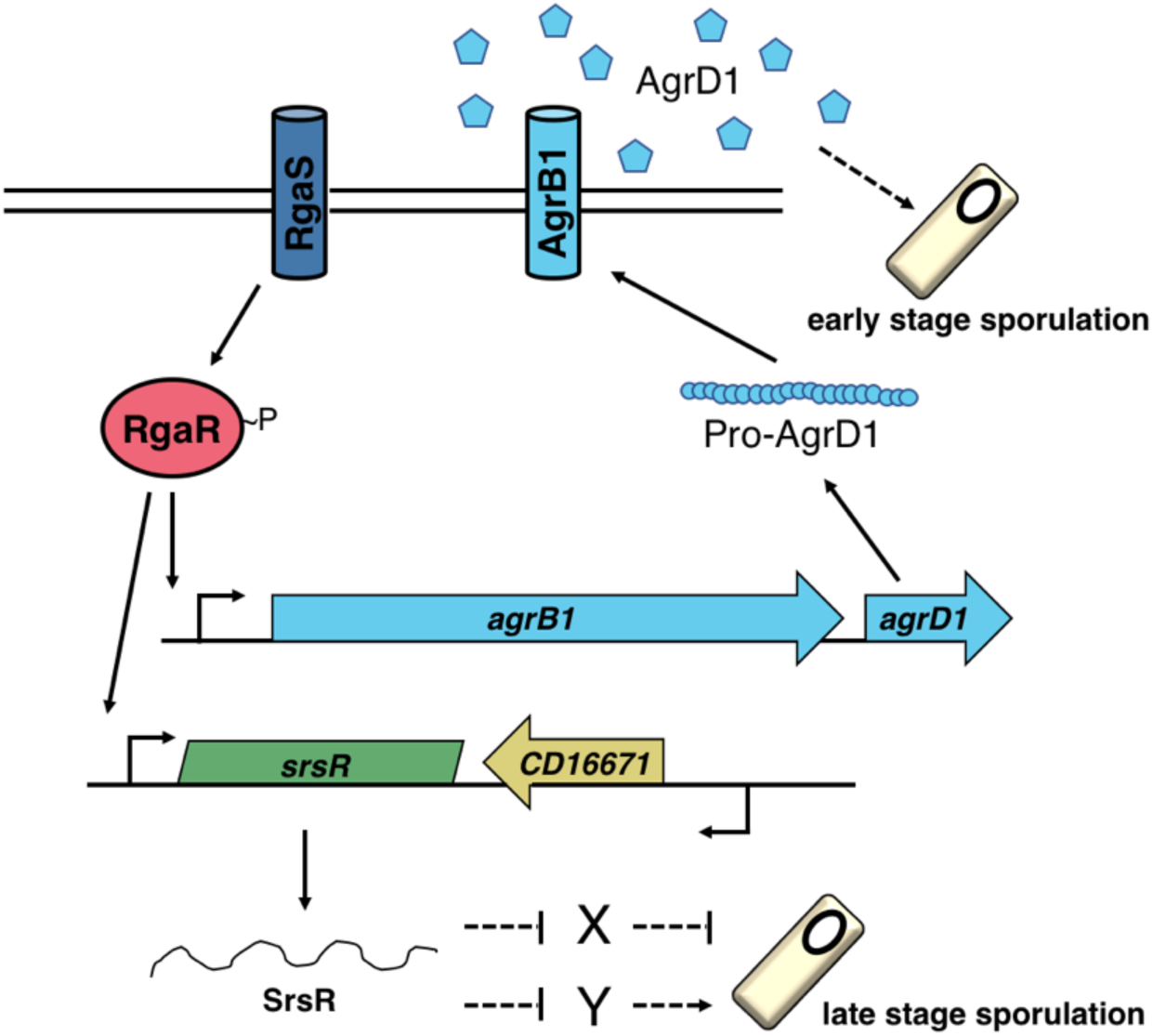
The model of RgaS-RgaR influence on *C. difficile* sporulation. The transmembrane sensor histidine kinase RgaS autophosphorylates in response to an unknown stimulus and transfers the phosphoryl group to the cytosolic response regulator RgaR. RgaR∼P activates transcription of its regulon, including *agrB1D1* and *srsR*. The gene products of the *agrB1D1* locus cleave and export the extracellular quorum-sensing peptide, AgrD1, which promotes early stage sporulation through an unknown factor. *srsR* encodes a small regulatory RNA whose abundance differentially impacts late stage sporulation via unidentified targets. Known direct interactions are depicted with solid lines while unknown regulatory pathways are denoted with dashed lines.

## METHODS

### Bacterial strains and growth conditions

The bacterial strains and plasmids constructed and used in this study are listed in **Table S2**. Plasmid construction details are in **Table S3**. *C. difficile* was routinely cultured in a Coy anaerobic chamber at 37°C in an atmosphere of 10% H_2_, 5% CO_2_, and balanced with N_2_, as previously described and grown in brain-heart infusion (BHI) broth supplemented with yeast extract (69–71). For plasmid maintenance or selection in *C. difficile*, 2-10 μg/ml thiamphenicol, 100-500 μg/ml spectinomycin, or 2-5 μg/ml erythromycin were supplemented in the media as needed. Overnight cultures were supplemented with 0.1% taurocholate to allow present spores to germinate and 0.2% fructose to prevent sporulation within the culture, as indicated (70,72). *Escherichia coli* were grown at 37°C in LB medium supplemented with 100 μg/ml ampicillin, 100 μg/ml spectinomycin, 300 μg/ml erythromycin, and/or 20 μg/ml chloramphenicol as needed, and *Bacillus subtilis* were grown at 37°C in LB (73) supplemented with 100 μg/ml spectinomycin or 1 μg/ml erythromycin as needed. *B. subtilis* was also supplemented with 3.5 mM KNO_3_ to support anaerobic growth. Kanamycin (100 μg/ml) was used to counterselect against *E. coli* and *B. subtilis* after conjugation with *C. difficile* (74).

### Strain construction

*C. difficile* 630 (GenBank accession no. NC_009089.1) and R20291 (GenBank accession no. NC_013316.1) were used as the templates for primer design, and *C. difficile* 630Δ*erm* and R20291 genomic DNA were used as the template for PCR amplification and mutant construction. The *C. difficile* 630Δ*erm* Δ*rgaS*, 630Δ*erm* Δ*rgaR*, 630Δ*erm* Δ*rgaS* Δ*rgaR*, 630Δ*erm* Δ*srsR-CD16671*, 630Δ*erm* Δ*CD16671*, 630Δ*erm* Δ*agrB1D1*, R20291 Δ*rgaS*, and R20291 Δ*rgaR* strains were constructed using allele-coupled exchange using either the pMSR or pMSR0 pseudo-suicide vectors for the 630 and R20291 backgrounds, respectively (41). Briefly, pMSR/pMSR0 vectors containing the targeted gene’s 5ʹ and 3ʹ homology arms, with a selective antibiotic resistance cassette in between, were conjugated into the respective parent strain and were selected for growth on 10 μg/ml thiamphenicol and 100 μg/ml kanamycin. Colonies were then passaged to either 500 μg/ml spectinomycin or 5 μg/ml erythromycin and subsequently grown in 10 ml BHIS with the selective antibiotic and 100 μg/ml anhydrotetracycline (ATC) to induce the double recombination event. After ATC treatment, isolated colonies were screened for successful allelic exchange by PCR. Complementation strains for the *rgaS* and *rgaR* mutants were constructed by either expressing the wild-type or site-directed mutated *rgaS* or *rgaR* alleles via the conjugative transposon Tn*916*. *rgaR* mutants expressing either P*cprA-agrB1D1* or P*cprA-srsR- CD16671* and the R20291 strain expressing P*cprA-srsR-CD16671* were also constructed using the conjugative transposon Tn*916*. All strains were confirmed by PCR analysis, and most were further confirmed by whole genome sequencing (SeqCenter, Pittsburgh, PA; see below).

The Benchling CRISPR Guide RNA Design tool was used to create sgRNAs targeting *rgaS*, *rgaR*, *CD0587*, *CD2098*, *CD15111*, *srsR*, and *agrB1* (35), which were generated by PCR and subsequently cloned into pMC1123. The details of vector construction are given in the supplemental material (see **Fig. S7**). Plasmids pMC1249 and pMC1250 were synthesized by Genscript (Piscataway, NJ).

### Illumina Whole Genome Sequencing

*C. difficile* strains were grown overnight in and harvested from 10 ml BHIS. After storing cell pellets overnight at -80°C, genomic DNA was isolated using a modified Bust ‘N’ Grab protocol (26,75). Contaminating RNA was removed by incubation with RNase A (Ambion) for 1 h at 37°C. Library prep and Illumina sequencing were performed by SeqCenter (Pittsburgh, PA). Briefly, sample libraries were prepared using the Illumina DNA Prep Kit and IDT 10bp UDI indices, and sequenced on an Illumina NextSeq 2000, producing 2x151bp reads. Demultiplexing, quality control, and adapter trimming was performed with bcl-convert (v3.9.3; Illumina; https://support-docs.illumina.com/SW/BCL_Convert/Content/SW/FrontPages/BCL_Convert.htm). Using Geneious Prime v2022.3, the resulting reads were paired and trimmed using the BBDuk plug-in and subsequently mapped to the reference genome (630Δ*erm*; NC_009089.1). The Bowtie2 plug-in was used to search for the presence of SNPs and InDels under default settings with a minimum variant frequency set at 0.85. Genome sequence files were deposited to the NCBI Sequence Read Archive (SRA) BioProject PRJNA986905.

### Sporulation assays and phase-contrast microscopy

*C. difficile* cultures were grown overnight in BHIS medium supplemented with 0.1% taurocholate, to promote germination of existing spores, and 0.2% fructose, to prevent sporulation during growth (70). *C. difficile* cultures were subsequently diluted with fresh BHIS, grown to mid-exponential phase (OD_600_ ≅ 0.5), and plated onto 70:30 agar supplemented with 2 μg/ml thiamphenicol and 1 μg/ml nisin as needed. After 24 h (H_24_), ethanol-resistant sporulation assays were performed as previously described (14,76). Briefly, cells were scraped from 70:30 agar and suspended in BHIS medium to an OD_600_ ≅ 1.0. Total vegetative cells per milliliter were determined by immediately serially diluting suspended cells in BHIS, plating onto BHIS plates, and enumerating colony forming units (CFU) after at least 36 h of growth. Simultaneously, spores per milliliter were enumerated after a 15 min incubation in 28.5% ethanol by serially diluting in 1X PBS with 0.1% taurocholate, plating onto BHIS with 0.1% taurocholate, and counting after at least 36 h of growth. The sporulation frequency was calculated as the total number of spores divided by all viable cells (spores plus vegetative cells). A *spo0A* mutant was used as a negative sporulation control. Statistical significance was determined using a one-way ANOVA, followed by a Dunnett’s or Tukey’s multiple-comparison test, as indicated (GraphPad Prism 9.5.0). Phase-contrast microscopy was performed H_24_, using the suspended cells from 70:30 agar, with a Ph3 oil immersion objective on a Nikon Eclipse Ci-L microscope. At least two fields of view were captured with a Ds-Fi2 camera. Results represent a minimum of three independent biological experiments.

### Enzyme-linked immunosorbent assay (ELISA)

*C. difficile* TcdA and TcdB from culture supernatants was quantified after 24 h growth of *C. difficile* strains in TY, pH 7.4, as previously described (77). Supernatants, diluted in the provided dilution buffer, were assayed in technical duplicates using the tgcBIOMICS kit for simultaneous detection of both TcdA and TcdB, according to the manufacturer’s instructions. The results represent the means and standard error of the means of four independent biological replicates, and statistical significance was assessed via a one-way ANOVA, followed by a Dunnett’s multiple-comparison test (GraphPad Prism 9.5.0).

### Quantitative reverse transcription PCR analysis (qRT-PCR)

*C. difficile* strains were cultured on 70:30 agar as described above, and cells were harvested at H_12_ (defined as 12 h after the cultures are applied to the plate) into 6 ml of 1:1:2 ethanol:acetone:dH_2_O solution and stored at -80°C. RNA was isolated and Dnase-I treated (Ambion), and cDNA was synthesized (Bioline) using random hexamers as previously described (16,18,78). qRT-PCR analysis was performed in triplicate on 50 ng of cDNA using the SensiFAST SYBR & Fluorescein kit (Bioline) with a Roche Lightcycler 96. The results were analyzed by the comparative cycle threshold method (79), using the *rpoC* transcript as the normalizer, and are presented as the means and standard error of the means for a minimum of three independent biological replicates. An one-way ANOVA, followed by a Student’s *t*-test or a Dunnett’s multiple-comparison test as applicable (GraphPad Prism v9.5.0), was used to assess statistical significance.

### RNA sequencing (RNA-seq) analysis

*C. difficile* strains were cultured on 70:30 sporulation agar as described above and harvested at H_12_ into 6 ml of 1:1:2 ethanol:acetone:dH_2_O solution and stored at -80°C. RNA was isolated and Dnase-I treated (Ambion). Samples were sent to Microbial Genomics Sequencing Center (MiGS; Pittsburgh, PA) where library preparation was performed using Illumina’s Stranded Total RNA prep Ligation with Ribo-Zero Plus kit and 10bp IDT for Illumina indices. Sequencing was done on a NextSeq200 giving 2x50bp reads. Demultiplexing, quality control, and adapter trimming was performed with bcl-convert (v3.9.3; Illumina; see reference above). Using Geneious Prime v2022.2.2, the reads were mapped to the reference genome (630Δ*erm*; NC_009089.1). The expression levels were calculated and then subsequently compared using DESeq2 (80).

### Alkaline phosphatase activity assays

*C. difficile* strains were cultured on 70:30 sporulation agar as described above and harvested at H_8_ (defined as eight hours after the cultures are applied to the plates). Alkaline phosphatase (AP) assays were performed as previously described (81) excepting the exclusion of chloroform. Technical duplicates were averaged and results represent the means and standard error of the means of three independent biological replicates. An one-way ANOVA, followed by a Tukey’s multiple-comparison test (GraphPad Prism v9.5.0), was used to assess statistical significance.

## ACKNOWLEDGEMENTS

We are grateful to the members of the McBride lab for their helpful suggestions, comments, and discussions throughout the course of this study. We are also thankful to Craig Ellermeier for the gift of pIA33. This research was supported by the U.S. National Institutes of Health through research grants AI116933 and AI156052 to S.M.M. The content of this manuscript is solely the responsibility of the authors and does not necessarily reflect the official views of the National Institutes of Health.

## SUPPORTING INFORMATION

**Figure S1. CRISPRi knockdown of *rgaS* and *rgaR* decreases *rgaS* and *rgaR* transcript levels.** Transcript levels of *rgaS* (**A**) and *rgaR* (**B**) at H_12_ in 630Δ*erm* psgRNA- neg (MC2065), 630Δ*erm* psgRNA-*rgaS* (MC2066), and 630Δ*erm* psgRNA-*rgaR* (MC2227) grown on 70:30 agar supplemented with 2 μg/ml thiamphenicol and 1 μg/m nisin, as indicated. The means and standard error of the means of at least three independent biological replicates are shown. *, P < 0.01 by a Student’s *t-*test.

**Figure S2. PCR confirmation of *rgaS* and *rgaR* mutants.** PCR verification of allelic replacement of *rgaS* (*CD0576*) and *rgaR* (*CD3255*) with the spectinomycin (*aad9*) or erythromycin (*ermB*) cassette, respectively in 630Δ*erm*, 630Δ*erm rgaS* (MC2228), and 620Δ*erm rgaR* (MC2229). Expected PCR products’ sizes are: 3319 bp for the wildtype *rgaS* allele and 3078 bp for the Δ*rgaS*::*aad9* allele (primers oMC3123/3124); 2668 bp for the wildtype *rgaR* allele and 3436 bp for the Δ*rgaR*::*ermB* allele (primers oMC3261/3262).

**Figure S3. Toxin production is unaffected in *C. difficile rgaS* and *rgaR* mutants.** ELISA of TcdA and TcdB present in the supernatant of 630Δ*erm*, 630Δ*erm* Δ*rgaS* (MC2228), and 630Δ*erm* Δ*rgaR* (MC2229) grown in TY medium, pH 7.4, at H_24_. The means and standard error of the means of four independent biological replicates are shown. The statistical analysis used for these data sets was a one-way ANOVA followed by Dunnett’s multiple comparisons test.

**Figure S4. CRISPRi knockdown of RgaR-dependent genes specifically and significantly decreases their transcript levels.** Transcript levels of *CD0587*, *CD2098*, *CD15111*, and *n00620* at H_12_ in 630Δ*erm* expressing psgRNA-*neg* (MC2065) or CRISPRi targets for *CD0587* (MC2269), *CD2098* (MC2270), *CD15111* (MC2271), or *srsR* (MC2272) grown on 70:30 agar supplemented with 2 μg/ml thiamphenicol and 1 μg/m nisin. The means and standard deviations of three independent biological replicates are shown. *, P < 0.0001 by a Student’s *t-*test.

**Figure S5. PCR confirmation of *srsR-CD16671* and *CD16671* mutants and *srsR* transcript length.** (**A, B**) PCR verification of allelic replacement of the *srsR-CD16671* locus with the spectinomycin (*aad9*) cassette in 630Δ*erm,* 630Δ*erm* Δ*srsR-CD16671* (MC2351; **A**) and 630Δ*erm* Δ*CD16671* (MC2363; **B**). Expected PCR products’ sizes are: 2165 bp for the wildtype *srsR*-*CD16671* allele, 2745 bp for the Δ*srsR-CD16671*::*aad9* allele, and 3033 bp for the Δ*CD16671*::*aad9* allele (primers oMC3457/3458). (**C**) Primers spanning the regions within the *srsR-CD16671* locus depicted in (**D**), were used to amplify cDNA and RNA (minus reverse transcriptase negative control; -RT) from 630Δ*erm* grown on 70:30 agar at H_12_ and 630Δ*erm* genomic DNA (gDNA).

**Figure S6. PCR confirmation of 630Δ*erm agrB1D1* mutants and R20291 *CDR20291_0503* (*rgaS*) and *CDR20291_3113* mutants *(rgaR*).** (**A**) PCR verification of allelic replacement of the *agrB1D1* locus with the spectinomycin cassette (*aad9*) in 630Δ*erm* and 630Δ*erm* Δ*agrB1D1* Expected PCR sizes are 2585 bp for the wild-type allele and 2930 bp for the *agrB1D1*::*aad9* allele (primers oMC3360/3361). (**B**) PCR verification of allelic replacement of *rgaS* and *rgaR* with the spectinomycin (*aad9*) cassette in R20291, R20291 Δ*rgaS* (MC2378), and R20291 Δ*rgaR* (MC2379). Expected PCR products’ sizes are: 3215 bp for the wildtype *rgaS* allele and 2974 bp for the Δ*rgaS*::*aad9* allele (primers oMC3124/3554); 2667 bp for the wildtype *rgaR* allele and 3056 bp for the Δ*rgaR*::*aad9* allele (primers oMC3261/3262).

**Figure S7. DNA cloning and vector details.**

**Table S1.** RNA-seq of *rgaR* mutant compared to 630Δ*erm*.

**Table S2. Bacterial strains and plasmids.**

**Table S3. Oligonucleotides.**

